# Understanding Harmonic Structures Through Instantaneous Frequency

**DOI:** 10.1101/2021.12.21.473676

**Authors:** Marco S. Fabus, Mark W. Woolrich, Catherine W. Warnaby, Andrew J. Quinn

## Abstract

The analysis of harmonics and non-sinusoidal waveform shape in neurophysiological data is growing in importance. However, a precise definition of what constitutes a harmonic is lacking. In this paper, we propose a rigorous definition of when to consider signals to be in a harmonic relationship based on an integer frequency ratio, constant phase, and a well-defined joint instantaneous frequency. We show this definition is linked to extrema counting and Empirical Mode Decomposition (EMD). We explore the mathematics of our definition and link it to results from analytic number theory. This naturally leads to us to define two classes of harmonic structures, termed strong and weak, with different extrema behaviour. We validate our framework using both simulations and real data. Specifically, we look at the harmonics structure in the FitzHugh-Nagumo model and the non-sinusoidal hippocampal theta oscillation in rat local field potential data. We further discuss how our definition helps to address mode splitting in EMD. A clear understanding of when harmonics are present in signals will enable a deeper understanding of the functional and clinical roles of non-sinusoidal neural oscillations.

## I. INTRODUCTION

NEUROPHYSIOLOGICAL recordings often show non-sinusoidal features [1]. Non-sinusoidality has been observed across multiple species and different modalities, and such features have functional roles [2], [3], [4] Non-sinusoidal waveforms have harmonics in their spectra, which can produce spurious results when using cross-frequency connectivity metrics, such as phase-amplitude coupling (PAC) [5], [6] and phase-phase coupling (PPC) [7]. This difficulty in distinguishing whether a signal comprises a single non-sinusoidal oscillation or several interacting oscillations has practical consequences for signal processing. For instance, harmonic coupling may account for most, if not all, PAC detected in human magnetoencephalography (MEG) studies [8].

To tackle this ambiguity, we need a complete definition for the question “what exactly is a harmonic?”. The reader might think this a trivial question, and agree with the definition given by Wikipedia: “A harmonic is a wave with a frequency that is a positive integer multiple of the frequency of the original wave, known as the fundamental frequency [9].” Existing literature in neuroscience often uses this ‘integer frequency ratio’ definition of a harmonic [10], with some authors including a stable phase relationship between harmonics as a condition [8]. We contend that these definitions are correct but incomplete in that they allow for a wide set of cases where intuitively separate signals would be labelled as harmonics. For instance, with one set of amplitude and phase values, the sum of a 10Hz and a 20Hz oscillation may create a single non-sinusoidal oscillation in which the 20Hz signal “blends in” to the 10Hz base (Fig. 1A). Another amplitude and phase configuration may create a summed signal in which dynamics from both the 10Hz and 20Hz signals are clearly and separately visible (particularly if the amplitude of the 20Hz signal is relatively high, Fig. 1B). The integer frequency ratio and consistent phase conditions are not sufficient to separate these cases.

**Fig. 1.**
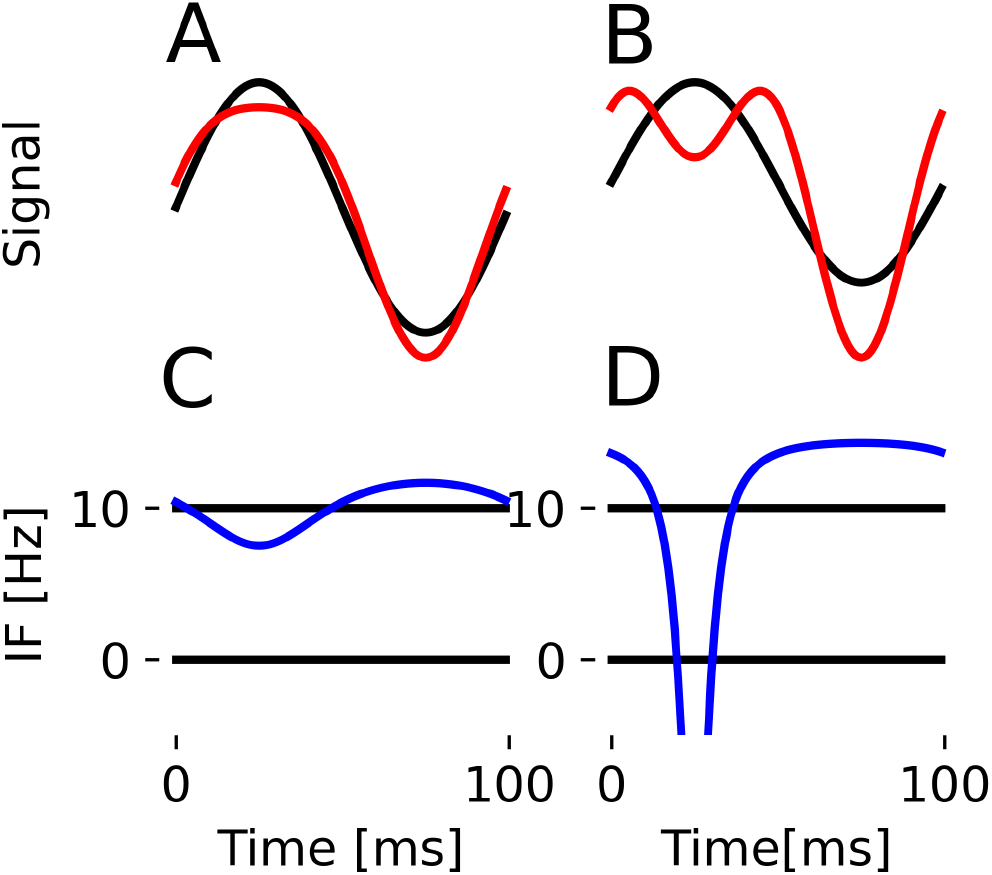
Harmonic intuitions. Top plots show the sum of a 10Hz wave (base, unit amplitude) and a 20Hz wave (HF) in red and a reference sine wave (black). In (A), the HF amplitude is low (0.2) and shape changes without introducing prominent extrema. In (B), HF amplitude is high (0.75) and new extrema are introduced. (C) and (D) show the associated instantaneous frequencies. (C) is well-defined everywhere, whilst (D) goes negative due to prominent secondary extrema.

How do we distinguish non-sinusoidality driven harmonics from independent oscillations (perhaps synchronised in frequency and coherent for purposes of information transfer [11] [12])? A complete and intuitive definition of when signals are in a harmonic relationship is lacking.

Empirical Mode Decomposition (EMD) is an alternative spectral decomposition method to Fourier-based techniques [13]. It decomposes data into a handful of Intrinsic Mode Functions (IMFs). EMD does this by a sifting algorithm where progressively slower oscillations are identified by finding extrema in the time-series. As such, it is intimately linked to the harmonic definition we propose. For instance, in a signal made of two sinusoidal components, EMD treats them as separate or joint based on which tone dominates the extrema rate [14]. Moreover, each IMF has a well-defined, non-negative Instantaneous Frequency (IF), theoretically able to represent any non-sinusoidal oscillation. However, in noisy real-world data, the bandwidth of an IMF is limited, even with improved EMD-based techniques [15]. Highly non-sinusoidal waveforms may thus have harmonics split across different IMFs, which we refer to as *mode splitting*.

Following intuitions from the Empirical Model Decomposition [13] [14], we propose that the Instantaneous Frequency [16] is the missing ingredient for a full definition of a harmonic. IF is able to fully characterise the shape of a mono-component, non-sinusoidal oscillation but will collapse into non-physical negative frequencies when representing a multi-component signal [2]. We can utilise this property to define an intuitive set of conditions for deciding when two signal components are in harmonic relation.

In this work, we propose a simple yet powerful set of conditions to define harmonics. We formalise the notion using instantaneous frequency and show this can be intuitively interpreted through notions of extrema counting. We find a natural interpretation of our results in the language of EMD. Choosing an analytically tractable model, we further explore the mathematical properties of our definition. We link them to results from analytic number theory and find two types of harmonics differing in their extrema. We then study harmonics in a simulated neuronal oscillations using the FitzHugh-Nagumo model. Finally, we apply our framework together with masked EMD to study rat local field potential (LFP) data. We validate our conditions on the asymmetric theta oscillation shape and illustrate how to decide whether to combine IMFs to address the mode splitting problem.

## II. HARMONIC STRUCTURES

### A. Intuition

What do we mean when we say oscillatory time-series are in a harmonic relation? In lay terms, we mean that one time-series, the *base*, determines “most” of the wave properties (e.g. the period, most of the amplitude), whereas the other time-series, the *harmonics*, determine fine details of the waveform shape. For an example, see Fig. 1. In top panels A and B, we see the sums of two waveforms, a base 10Hz sine and a 20Hz cosine. In panel A, the 10Hz waveform is five times the amplitude of the 20Hz waveform. The waveforms have an integer frequency ratio and a constant phase relationship, which guarantees the resulting waveform has the same period as the base sine function. Additionally, the joint waveform has a well-defined instantaneous frequency (panel C). Following from the Introduction, harmonic signals thus blend into a single waveform. In panel B, the 20Hz waveform is now 0.75x the amplitude of the 10Hz waveform. New prominent extrema appear in each resulting cycle. The summed signals retain their dynamics, making them nonharmonic. Instantaneous Frequency (IF) is emerging as a robust way to characterise non-sinusoidal waveform shape, but this frequency only makes sense if it is non-negative [2]. Prominent extrema are what causes IF to be negative, as in panel D. As such, we propose to define that in addition to an integer frequency ratio and a constant phase relationship, harmonics are signals that added to the base have a non-negative joint IF. The reader is encouraged to further explore the link between harmonics and instantaneous frequency using our custom shape generator and an interactive notebook accompanying this paper. The latter is a repository which includes code to reproduce all figures in this paper.

### B. Formalising Harmonic Conditions

Here we formalise the above intuitions. We shall say that the resultant signal *x*(*t*) formed as a sum of *N* sinusoids ordered by increasing frequency,

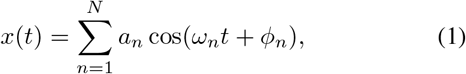

is to be considered a harmonic structure if:

1. All sinusoids have an integer frequency relationship to the base and a constant phase relationship, i.e. *ω_n_* = *nω*_o_, *n* ∈ ℤ and 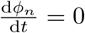,
2. The joint instantaneous frequency *f_J_* is finite and non-negative for all *t*, i.e. (*f_J_* ≥ 0)∀*t*, where *a_n_* are the sinusoids’ amplitudes.

The first condition is the same as that typically used in the literature [8]. In lay terms, it means the joint waveform repeats and is the same at *t* and *t*+*T*, where *T* is the period of the base function. We can easily show this: Noting that *ω*_1_ = 2*π/T*,

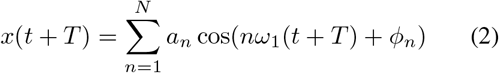

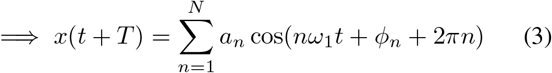

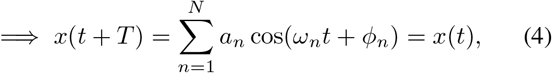

as all functions are 2*π* periodic.

Our signal model considers continuous oscillations with constant *a_n_*, but we note that all of the analysis in this paper also applies if this waveform experiences amplitude modulation (so long it happens slowly with a time scale *T_AM_ >* 2*π/ω*_1_). This is a consequence of Bedrosian’s Theorem [17] and is outlined further in the Discussion.

### C. Instantaneous Frequency

We aim to understand non-sinusoidal signals through instantaneous frequency, which fully characterises waveform shapes.

Hence, in this section, we analytically derive the instantaneous frequency for our signal model (1).

Following [13], we define the instantaneous frequency using the analytic signal phase. For a real signal *u*(*t*), define its analytic counterpart as *x_A_* = *u*(*t*) + *iv*(*t*), where *v*(*t*) is the Hilbert transform of *u*(*t*). We can rewrite the analytic signal as *x_A_* = *A*(*t*)*e^iθ^*^(*t*)^, where the instantaneous phase is obtained from the real and imaginary components of *x_A_* as tan *θ* = *v/u*. From this we define the instantaneous frequency as

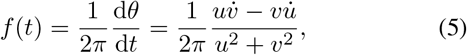

where 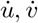 signify the time derivatives. This right-most expression is derived in Appendix A.

Using linearity of the Hilbert transform and equation (5), we can find the general joint instantaneous frequency for our sum of sinusoids (1):

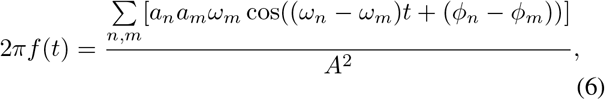

where the denominator is

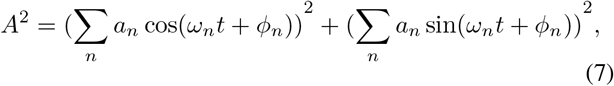

and we used the standard Hilbert transform *v* = *a_n_* sin(*ω_n_t* + *ϕ_n_*) for *u* = *a_n_* cos(*ω_n_t* + *ϕ_n_*). The full derivation of this expression is in Appendix B.

### D. *Case N* = 2 *and Link to Extrema Counting*

Harmonic condition 2 requires the instantaneous frequency to be non-negative. Recall that this is based on the premise that signals that cause prominent extrema are not harmonics, and that prominent extrema cause ill-defined, negative IF (see Fig. 1). Here we illustrate how harmonic condition 2 is linked extrema present in the waveform (Fig. 2).

**Fig. 2.**
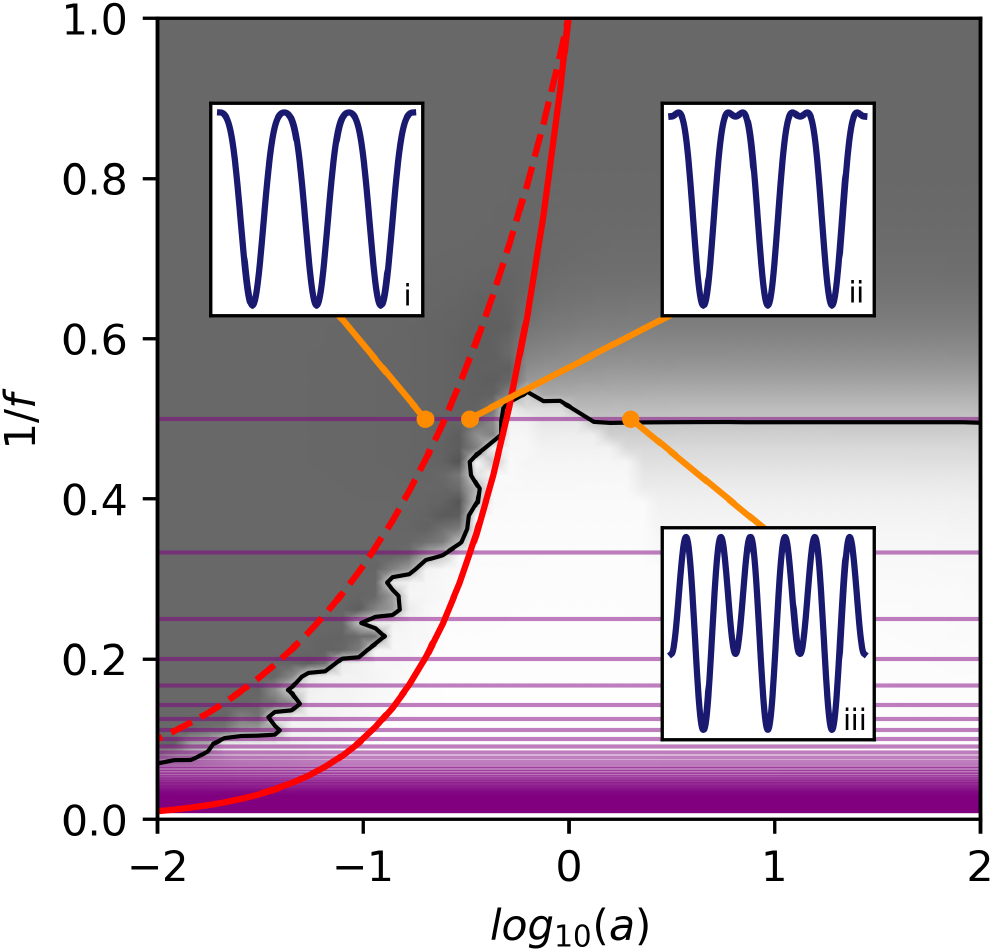
EMD separation of simulated two-tone signals with amplitude ratio *a* and frequency ratio *ω* in relation to harmonics. Gray shading shows whether EMD treats signals as separate oscillations (light) a single waveform (dark), or a mixture of waveforms (gray). Purple lines show where harmonic condition 1 (integer frequency ratio) is satisfied. The lines *aω* = 1 (red, solid) and *aω*^2^ (red, dashed) are also shown. Insets show three possible types of two-tone signals. (i) A strong harmonic structure - HF adds to the non-sinusoidal shape with no secondary extrema.(ii) A weak harmonic structure - small secondary extrema are present but the joint IF is still well-defined. (iii) Two tones are separate and not harmonically related. Strong secondary extrema are present and the IF is not well-defined. The separation map is reproduced from [14].

It is instructive to consider the case with *N* = 2. With only two sinusoids, we can write our signal model (1), without loss of generality as

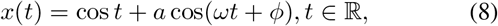

where *a* and *ω* are the amplitude / frequency ratios of the waveforms and *ϕ* their phase difference. We further restrain ourselves to the case *ω >* 1, such that the cos *t* term can be referred to as the base (or lower frequency component, LF), with the other termed the potential harmonic (or higher frequency component, HF). This simplified case follows that of [14], except for swapping *a* and *ω* (*f* in the original paper) into the HF term for greater clarity.

We now re-state the harmonic conditions for the case of *N* = 2, i.e. the conditions that need to be met for us to consider HF as a harmonic to LF.

The first condition simply amounts to *ω* = *n, n* ℤ (purple lines in Fig. 2), and *ϕ* = 0.

The second condition requires that the joint IF is non-negative for all time points. The joint IF from (6) simplifies to

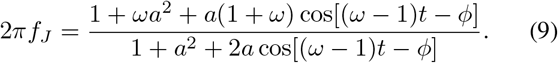

The denominator in (6) is always non-negative, so to have a non-negative joint IF we demand a non-negative numerator, noting the minimum value of a cosine expression is −1:

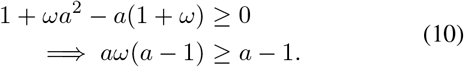

If *a >* 1, we can divide both sides by (*a* − 1) freely, but if *a <* 1, we must flip the equality sign when dividing by (*a* − 1). Case *a* = 1 satisfies the inequality trivially, thus we obtain the following restrictions on *a* and *ω*:

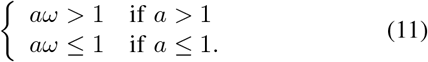

As was demonstrated in [14], the *aω* multiple is a key determinant of extrema locations in a joint two-tone signal such as *x*. Specifically, *aω >* 1 means the extrema rate is exactly the same as that of the HF component. HF is the dominant mode, and we could consider LF to be its ‘sub-harmonic’. However, harmonics typically have progressively lower amplitudes (*a <* 1), so we shall reserve the term ‘harmonic’ only for the cases of decreasing amplitudes. This case might potentially be useful when fixing specific instances of EMD mode splitting issues.

If *a* ≤ 1, in order to have a well-defined IF, we demand the extrema rate to *not* be set by HF. If *aω*^2^ *<* 1, it is set by LF ([14], Fig. 2), otherwise the extrema rate is a mixture of HF and LF extrema. The *a <* 1 case is the more the common situation for non-sinusoidal functions present in neurophysiological recordings.

The *aω* = 1 line is also crucial in the behaviour of EMD when splitting signals [14]. In the gray shading of Fig. 2, we can see the space of potential harmonics crosses both regions where EMD treats tones separately and as one modulated signal. This means that waveform shape reconstruction (combining several IMFs) may be necessary in cases where EMD separates them. This issue is even more widespread in real-world data including noise due to dyadic behaviour intrinsic to EMD [18]. If two IMFs satisfy the harmonic conditions, we can say to have identified a base and a harmonic. Due to linearity of the Hilbert transform, adding them to produce a single broad instantaneous frequency shape is then valid.

Superimposed on the EMD separation map, Fig. 2 shows the possible types of joint two-tone signals. For low amplitudes such that *aω* ≤ 1, the joint waveform forms a harmonic structure as its properties are dominated by the LF base and its joint IF is well-defined (inset (i)). Small secondary extrema are present when *aω*^2^ > 1 (inset (ii)). This anticipates the distinction between strong and weak harmonics we describe below. Finally, if the HF amplitude is too high (inset (iii)), the IF ceases to be well-defined, large secondary extrema are present, and we no longer consider HF to be a harmonic.

We can also re-write this result in a more general form that will be useful when considering multiple harmonics. Because *ω* = *n*, we can write *aω* ≤ 1 ⟹ *an* ≤ 1 ⟹ *a* ≤ 1/*n*. Similarly, *aω*^2^ ≤ 1 ⟹ *a* ≤ 1/*n*^2^. Thus amplitude conditions are of the form *a* = 1/*n^γ^*, where *γ* is a real exponent. For *γ* ≥ 1, we are guaranteed a non-negative joint instantanous frequency.

In summary, the *N* = 2 case illustrates key insights into harmonic systems. We see how demanding the instantaneous frequency to be non-negative is directly linked to the presence of extrema and the extrema rate. This is linked to EMD as it is a decomposition technique built on sifting extrema. We also see that cases with non-negative IF may have small secondary extrema, and whether these are present anticipates the strong/weak harmonic types introduced below.

### E. Examples of Harmonic Structures

Here we briefly consider three common examples of periodic signals with strong harmonics - the triangular wave (*y*_1_), saw-tooth wave (*y*_2_), and square wave (*y*_3_). Electrophysiological data often shows aspects of these waves (e.g. the ‘flat top’ of motor mu waves [19]), so they serve as a useful reference point. Their Fourier series are well-known:

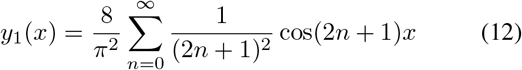

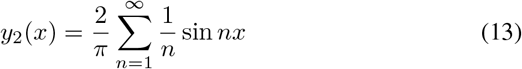

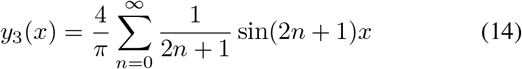

All three trivially satisfy Condition 1 as their frequency ratios are integers only and all phases are constant and zero. We are in the regime *a <* 1 for all three as HF harmonics get progressively smaller in amplitude. From the Fourier coefficients, the *a_i_ω_i_* product for neighbouring harmonics *n* and *n* + 1 in structure *y_i_* is as follows:

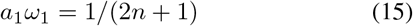

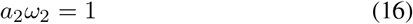

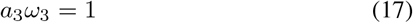

It is clear all three examples satisfy the *aω* ≤ 1 condition for neighbouring harmonics and are thus bona fide harmonic structures as expected. Alternatively, given the frequencies are some integers *m*, the amplitude falls as *a* = 1/*m^γ^* with *γ*_1_ = 2, *γ*_2,3_ = 1. These waveforms and their instantaneous frequencies are plotted in the Appendix Figure 8.

### F. Case N = 3

The general case of (6) is not conducive to simple conditions such as (11). However, we know that in reality, the amplitude falls with frequency. For this and the following section, we explore the amplitude-frequency relationship of the form *ω_n_* = *n*, *a_n_* = 1/*n^γ^*, that is *a_n_ω_n_* = 1/*n^γ−^*^1^ with *γ* ≥ 1 ∈ ℝ. This is the generalised form of the standard harmonic structures above and it will turn out to be insightful in the *N* = ∞ case. The reader can explore a wide variety of harmonic structures and their IF in the interactive notebook attached to this paper. The case of an exponentially falling amplitude is explored in Appendix E.

We have seen that for *N* = 2, *γ* ≥ 1 always leads to a non-negative instantaneous frequency. Here we show how this is modified in the case of *N* = 3. This is relevant e.g. for an EMD sift where an IMF includes two harmonics [7].

Our signal model is

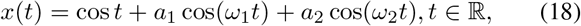

with *a_n_* = 1/*n^γ^*, *ω_n_* = *n*, and a constant phase assumed. We again use (6), noting its denominator is always positive. As such, demanding a non-negative IF to satisfy harmonic condition 2 means

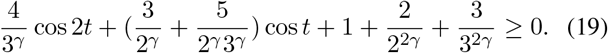

We can rewrite this as a quadratic in cos *t* using the double angle formula and compute the discriminant to find the restrictions on *γ*. This is done in Appendix D. Here we note that the critical exponent *γ_c_* which guarantees a non-negative IF is found as the solution to

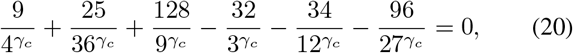

which we numerically find to be *γ_c_* = 1.0177, which is only very slightly different from the *N* = 2 case where *γ_c_* = 1. As the number of harmonics in a given IMF can be expected to be small, we can therefore apply the *aω* ≤ 1 condition to find harmonic structure.

### G. Case N = ∞ and Two Types of Harmonics Structures

In this section, we use an analytically tractable harmonic model to look at shapes with an infinite number of harmonics. In doing this, we find some shapes gain no secondary extrema even with infinitely many harmonics, whilst some do. We use this to classify harmonics into two types.

In the previous section, we found that for *N* = 3, requiring a non-negative IF is equivalent to having a critical exponent *γ >* 1 for a signal model (6) with *a_n_* = 1/*n^γ^* and *ω_n_* = *n*. Here we ask the question: if we have an infinite number of harmonics, are there any exponents *γ* for which no secondary extrema are introduced?

Consider the the sum to infinity of the numerator in (6). If all phases have the same value, the IF has a maximum at *t* = 0 and, where all cosines add constructively. For a well-defined waveform, IF needs to remain finite even in the infinite limit. If all phases are the same, consider the case with all *ϕ_n_* = 0 without loss of generality. The numerator becomes

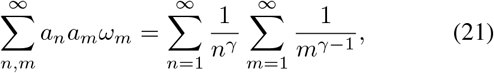

where as mentioned we again used the form *ω_n_* = *n*, *a_n_* = 1/*n^γ^*. We recognise these sums as the hyper-harmonic series (p-series). These diverge to infinity for exponent values ≤ 1 [20], and lie on the real line of the Riemann Zeta function for exponent values > 1 [21] [22]. Thus, including the denominator, we can write

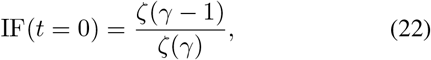

where 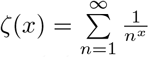. This converges to a finite real number only provided *γ >* 2. We therefore get two types of harmonic structures:

1. *Weak harmonic structures*. These have neighbouring harmonic that can be added to form a well-defined IF, but their IF diverges to infinity in the *N* = ∞ limit and ceases to be well-defined. They have *γ* ≤ 2, such as (13), and have small secondary extrema.
2. *Strong harmonic structures*. These have a well-defined (non-negative and finite) instantaneous frequency, even in the infinite limit. Harmonics do not introduce any new extrema and *γ >* 2. An example is

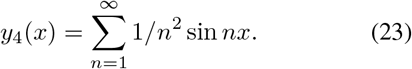

This distinction can be observed in Fig. 3. A weak harmonic structure (left) has small secondary extrema, whereas a strong harmonic structure (right) does not.

**Fig. 3.**
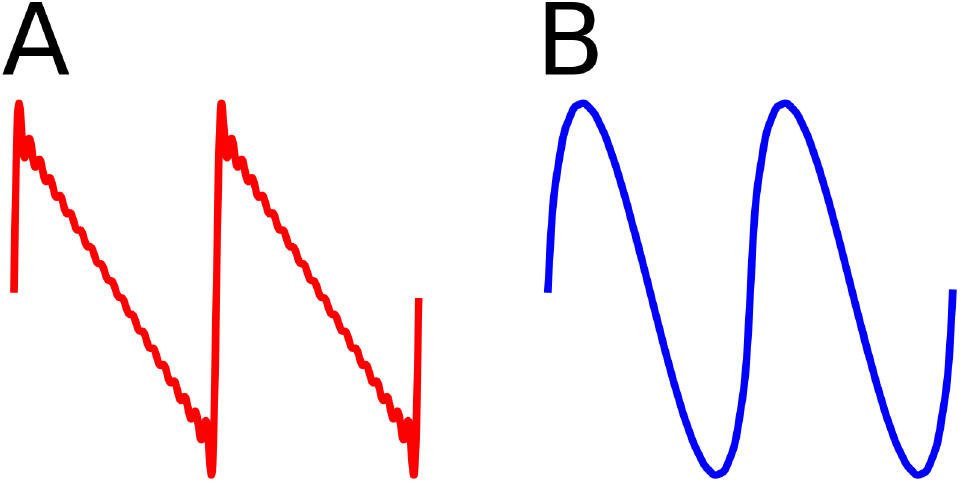
Two types of harmonic structures. (A) Weak harmonics: First *N* = 20 harmonics from (13). Secondary extrema are present and IF tends to a delta function. (B) Strong harmonics: First *N* = 20 harmonics from (23). No secondary extrema are present and the waveform is smooth with a finite IF in the *N* = *∞* limit.

To illustrate how no new extrema are present in strongly harmonic structures, we can consider an analytically tractable example of (1) with *a_n_* = 1/*n^γ^*, *ω_n_* = *n* and *ϕ_n_* = 0 (Fig. 5). Restrict ourselves to the case of even *N* for simplicity. These structures have an extremum at *t* = *π* for any number of harmonics. If this is to be the only extremum in (0, 2*π*), it must be convex. This is as a concave extremum would imply a local maximum and thus at least two additional secondary minima either side as the function must eventually turn to form maxima at *t* = 0 and *t* = 2*π*. We therefore demand a positive second derivative at *t* = *π* for no new secondary extrema:

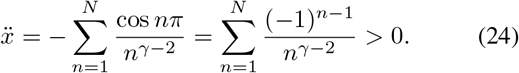

This sum as a function of *γ* is plotted in panel (C) of Fig. 5 for example values of *N*. We can see the second derivative is positive above *γ >* 2, indicating no new extrema are present as proposed. Interestingly, this function converges to the Dirichlet *η* function [23] in the *N* = limit, though its properties were not used here as we are interested in even *N* cases only. Odd *N* cases have an odd number of secondary extrema and are thus more tedious to analyse. We finally note here that this weak/strong distinction can be also understood as a constraint on the amplitude part of the analytic signal *x_A_* = *ae^iθ^* cardioid traced out in the complex (or equivalently polar) plane. This is illustrated in the middle panels of Fig. 5.

**Fig. 4.**
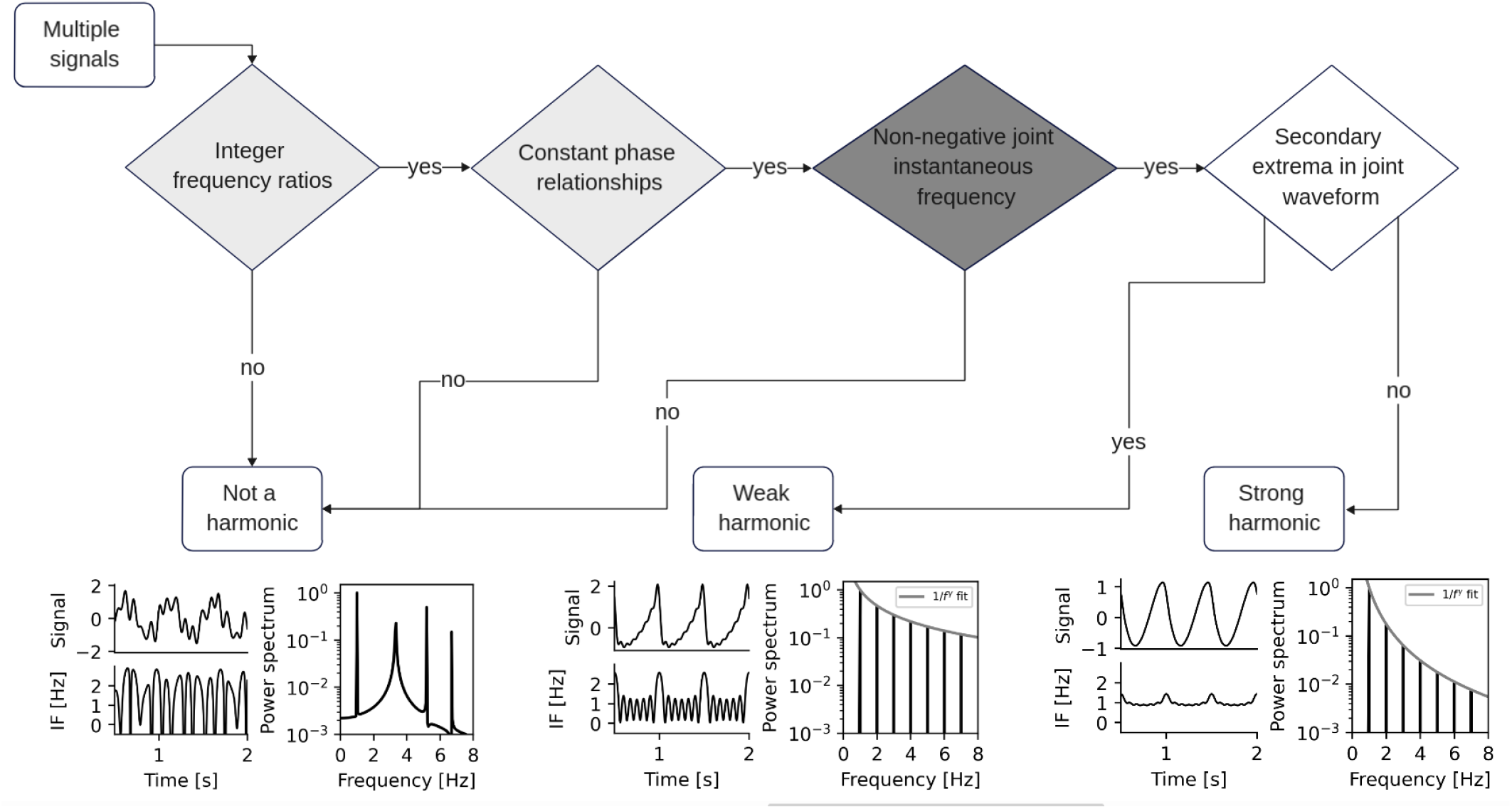
Harmonic assessment decision tree. Depending on whether signals meet condition 1 (light grey), condition 2 (dark grey), and how quickly harmonic amplitude falls, the joint signal is either not a harmonic (bottom left, multiple oscillatory processes present and IF sometimes negative), a weak harmonic structure (bottom centre, small secondary extrema present but IF non-negative), or a strong harmonic structure (bottom right, no secondary extrema).

**Fig. 5.**
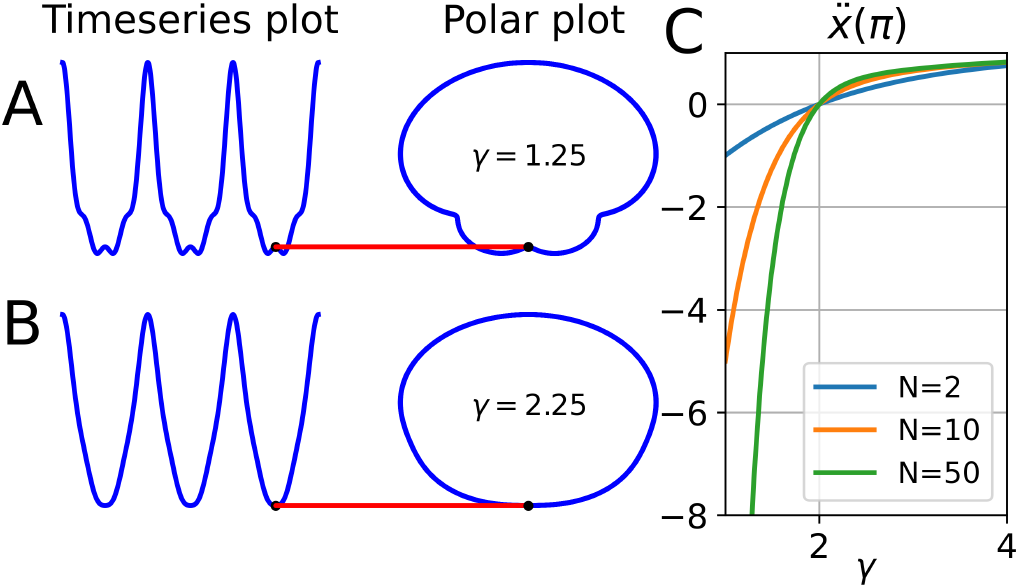
Two types of harmonics driven by the analytic amplitude term. (A) Weak harmonic structure with *γ* = 1.25. Left - time series (4 harmonics), right - equivalent polar plot of *ae^iθ^*. A secondary maximum is present at *t* = *π* (red line). (B) Strong harmonic structure with *γ* = 2.25. Left - time series (4 harmonics), right - equivalent polar plot of *ae^iθ^*. No secondary maximum is present as curves are convex at *t* = *π*. (C) Second derivative of a sum of cosines at *t* = *π* for different numbers of harmonics *N* It is clear the concave/convex transition is at *γ* = 2, marking the change from weak to strong harmonics. Note both curves (A) and (B) here have a well-defined IF > 0 everywhere.

In summary, a well-defined IF and a harmonic amplitudes falling off fast enough means harmonics introduce no new extrema, which we class as *strong* harmonics. Harmonic structures with amplitude modulation dynamics introducing small secondary extrema are then of the *weak* type. We studied the analytically tractable harmonic model of *a* = 1/*n^γ^* because it appears in common waveshapes. However, below we also present results from a more realistic neural waveshape based on the FitzHugh-Nagumo model and in the Appendix we look at exponentially decaying harmonic amplitudes.

### H. Summary: How to assess a harmonic

In this section, we summarise key metrics we have identified whilst exploring the theory of harmonic structures above. We list their role in studying harmonics and their practicability. Assessing harmonics using these quantities can be reduced to a simple decision tree (Fig. 4).

- *ω_n_*, the frequency ratio between signals. Needs to be an integer for harmonic structures. Tests whether signals align such that the base period is unchanged. For neurophysiological data, it is easy to check from peaks in the power spectrum.
- *ϕ_n_*, the phase relationship between signals. Needs to be constant for harmonic structures. Can be checked using metrics such as the phase locking value or distance correlation.
- *f_J_*, the joint instantaneous frequency. Non-negative if signals are harmonics. Tests that the waveform does not contain prominent extrema which would indicate separate oscillatory dynamics. It can be assessed in multiple ways:

– Equation 6 - analytical expression for when all parameters are known. Useful for forward modelling and simulation, but impractical to assess in real signals.
– *aω*, the harmonic ratio multiple. Tests a single pair of components with amplitude ratio *a* and frequency ratio *ω*. Computationally cheaper and easier to use than assessing *f_J_ >* 0 directly.
– The Hilbert transform. In complex signals, instantaneous frequency can be numerically estimated with software packages such as emd in Python [24].
- *γ*, the harmonic amplitude drop-off exponent. It distinguishes weak and strong harmonic structures, i.e. the presence of *any* secondary extrema. Assesses any number of harmonics at once. It is a good fit if amplitude roughly falls according to a power law.

## III. METHODS

Analysis of experimental data was done in Python 3.9.0. EMD was applied using the Python EMD package (v0.4.0), available with tutorials at https://emd.readthedocs.io/ [24]. Packages NumPy [25], SciPy [26], dcor [27], and Statsmodels [28] were used for analysis. Package Matplotlib [29] was used for plotting.

### A. Simulations

We simulated 10 seconds of a FitzHugh-Nagumo neuron with a sampling rate of 100kHz and parameters giving rise to a continuous 25Hz oscillation (stimulation current *I* = 0.475, initial membrane potential *V*_0_ = 0, recovery parameter *W*_0_ = − 0.4, scaling parameters *a* = 0.7, *b* = 0.8, and *τ* = 12.5 [30]). This is a dynamical system governed by the coupled equations

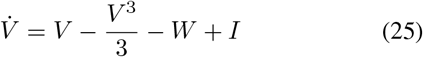

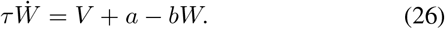

Its instantaneous frequency was computed with the Hilbert transform (emd.spectra.frequency transform) and its power spectrum with the Fourier transform (scipy.fft.fft). Harmonic peaks were found using scipy.signal.find peaks. For comparison with our analytical results, the log harmonic peak amplitudes were fitted against their log frequency using linear regression to estimate the harmonic amplitude drop off exponent *γ*.

### B. Rat Data

The rodent hippocampal theta oscillation is known to be non-sinusoidal [31], [32]. Therefore, to demonstrate our results on experimental data, we chose a publicly available hippocampal data set of Long-Evans rats [33], [34]. A 1000s local field potentials (LFP) recording from rat EC-013 sampled at 1250Hz was used for analysis. The electrode analysed was implanted in the hippocampal region CA1. The recording was first downsampled to 625Hz using scipy.signal.decimate. EMD sift was then computed with *N*_IMF_ = 8 modes using the mask sift [35] with the first mask frequency found from zero crossings in the signal and the rest as divided by increasing powers of 2. The sift threshold was 10^*−*8^ and mask amplitudes were all equal to the standard deviation of the input signal. Instantaneous phase, frequency, and amplitude were computed from the IMFs using the Hilbert transform with an instantaneous phase smoothing window of five time-points. The base theta IMF was chosen as that whose average instantaneous frequency was closest to the Fourier spectral theta peak estimated using Welch’s method (peak in 4-8Hz, function scipy.signal.welch, 8s segment length / 0.125Hz resolution). Individual cycles were computed from jumps in the wrapped instantaneous phase. To discard noisy cycles, only cycles with monotonic instantaneous phase and instantaneous amplitude above the 50th IA percentile were used for further analysis. Cycles were phase-aligned with *N* = 48 phase points and the shape was represented by the mean of the phase-aligned instantaneous frequency [2].

Next the harmonic conditions were tested. The recording was split into 20 segments of 50s each. Recall that the first condition is requires integer frequency ratios and a constant phase relationship between signals. Therefore, the first condition was taken to be satisfied if (i) the ratio between mean IF of HF mode and base was not significantly different from an integer, tested with a one-sample t-test with the nearest integer as the null hypothesis, and (ii) the base and HF had a constant phase relationship indicated by a significant distance correlation between their whole-recording instantaneous phases tested with the Student’s t-test. The distance correlation was used because it captures any statistical dependence between phases, not just a linear relationship (Pearson correlation) or a monotonic relationship (Spearman correlation). The second condition (that the joint IF is non-negative) was met if the amplitude and frequency ratios between HF and base satisfied the *aω* ≤ 1 condition outlined above. To classify the harmonic structure, the value of *aω*^2^ was also tested. We used the amplitude and frequency ratios instead of testing for non-negative IF directly because details of IF calculation are often unreliable due to noise and EMD sift issues.

## IV. RESULTS

### A. Simulations

We first explored harmonics in a simulated FitzHugh-Nagumo neuron spiking continuously at 25Hz (Fig. 6). The waveform was highly non-sinusoidal, as has been noted by other researchers [1]. This meant its instantaneous frequency trace ranged between 11Hz and 80Hz. Importantly, the IF always remained positive. Together with harmonic peaks in the power spectrum at integer multiples of 25Hz, their constant phase relationship, and an unchanged period, this meant the waveform indeed qualified as a harmonic structure in our framework.

**Fig. 6.**
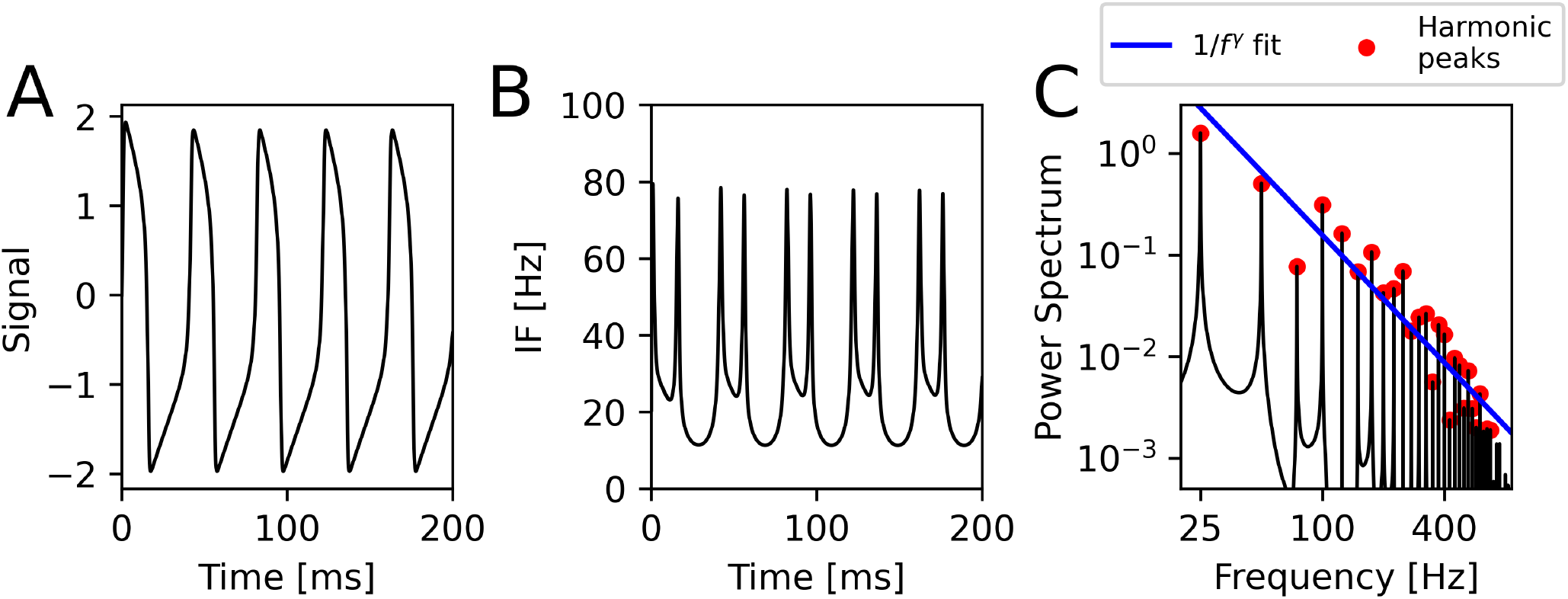
Harmonics in the FitzHugh-Nagumo neuronal spiking model. (A) 200 ms of membrane voltage of the neuron. Model parameters were chosen to produce continuous 25Hz spiking and the oscillation waveforms are highly non-sinusoidal. (B) Instantaneous frequency of the waveform in (A). Sharp edges correspond to higher frequencies. (C) Power spectrum of the data. A base at 25Hz with harmonics at each following 25Hz increment are clearly visible. Red dots signify individual harmonic peaks. Blue line is the linear fit to harmonic amplitudes in the log-log plane. The waveform is a strong harmonic structure (no secondary extrema) and harmonics fall off roughly as *an* = 1/*n*^2.08^.

In the theoretical section, we explored harmonics where amplitude falls as *a_n_* = 1/*n^γ^*. To compare our simulated neuron to this analytically tractable model, we performed a linear fit of the harmonic peak amplitudes in the log-log plane. Harmonic amplitude fell as approximately *a_n_* = *k/n*^2^.083 (Pearson *r* = −0.936, *P* = 3.69 × 10^*−*13^. This confirmed our analytical model was useful as an approximation to a system simulating the behaviour of neurons. Moreover, the waveform was a strong harmonic structure (it had no secondary extrema), just as we would predict with *γ >* 2.

### B. Rat Data

We validated our harmonic framework by applying it to the known non-sinusoidal hippocampal theta waveform. Masked EMD extracted four IMFs of interest (Fig. 7). IMF-4 was identified as the dominant base oscillation as it was closest in frequency to the Fourier theta peak. We then tested if IMFs higher in frequency showed a harmonic relationship with this base by applying the harmonic conditions (Tab. I).

**Fig. 7.**
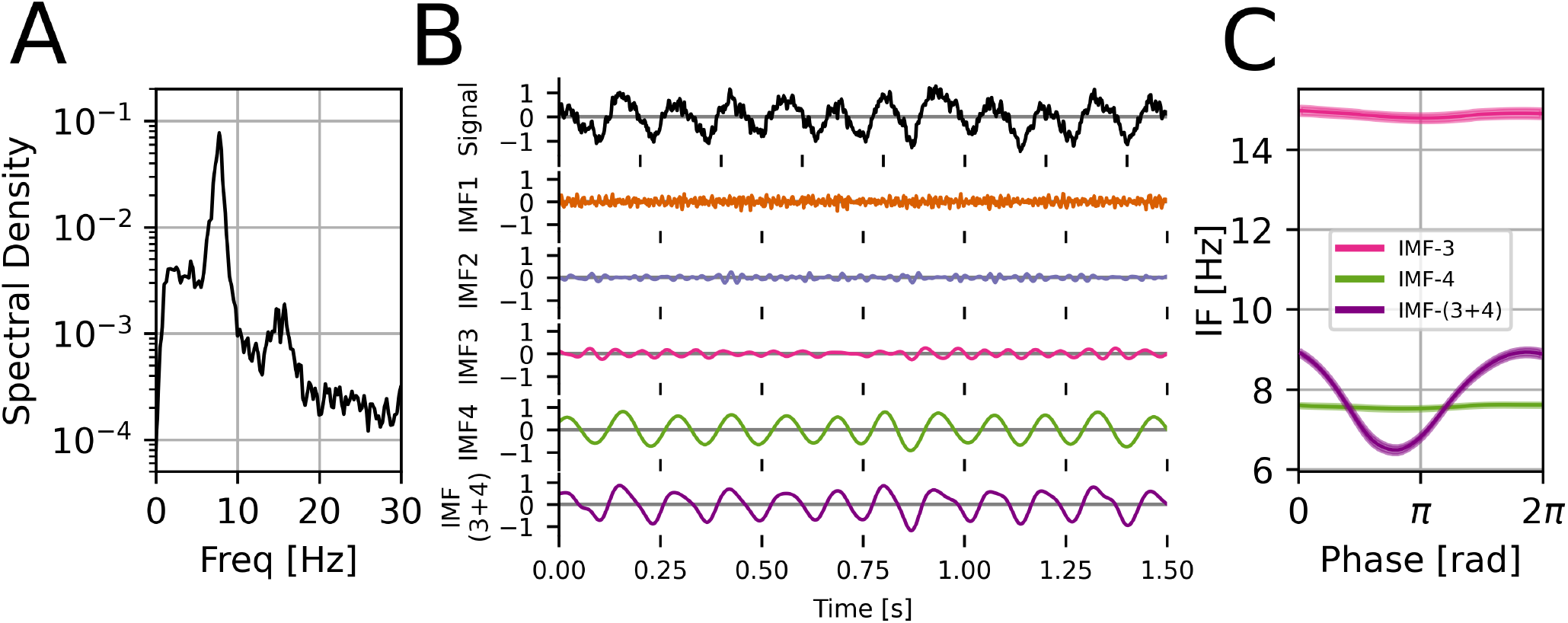
Harmonics in rat local field potential (LFP) data. (A) Power spectral density of the data. A base around 7.5Hz with a harmonic around 15Hz are clearly visible. (B) Example 1.5s of masked EMD sift results. Base is in IMF-4 and harmonic is in IMF-3 due to limited IMF bandwidth in the presence of noise. Joint waveform is shown as IMF-(3+4) (C) Phase-aligned IF (mean ± SEM across cycles). Both IMF-3 and IMF-4 are nearly sinusoidal. After verifying harmonic conditions between IMF-3 and IMF-4 are met, IMFs are added to reconstruct the full non-sinusoidal shape (purple).

Condition 1 is equivalent to testing for an integer frequency ratio and a stable phase relationship. IMF-1 and IMF-3 both showed a frequency ratio to the base not significantly different from an integer (11 and 2 respectively, P > 0.05). However, only IMF-3 also showed a significant (P *<* 0.001) and modest distance correlation (0.28) between its instantaneous phase and that of the base, showing presence of phase-phase coupling. IMF-3 had a very small (0.03), but also significant distance correlation.

Condition 2 was tested by the *aω* ≤ 1 relationship where *a* was the ratio of mean instantaneous amplitudes and *ω* the corresponding ratio for instantaneous frequencies. The *aω* products of IMF-2 and IMF-3 with the base signal were significantly below 1 (P = 1.7 × 10^*−*4^ and P = 2.9 × 10^*−*15^ respectively, one-sample Bonferroni-corrected t-test).

Taking both conditions into account, we found IMF-3 and IMF-4 to be robustly harmonically related. Each individual condition (integer frequency ratio, constant phase relationship, *aω* ≤ 1) was inconclusive on its own, showing the importance of a full definition of a harmonic.

Next we computed the mean of phase-aligned instantaneous frequency across cycles as a measure of waveform shape [2]. Both IMF-3 and IMF-4 showed nearly sinusoidal cycles. Taking their harmonic relationship and the linear nature of the

**TABLE 1.** RESULTS IN RAT LFP DATA

| IMF | IF [Hz] | IA [mV] | DCor | $\omega$ | $a$ |
| --- | --- | --- | --- | --- | --- |
| 1 | 81.8 (3.3) | 0.16 (0.02) | 0.007 | **10.9 (0.3)** | 0.27 (0.04) |
| 2 | 31.7 (1.0) | 0.11 (0.01) | **0.030** | 4.2 (0.2) | **0.18 (0.04)** |
| 3 | 14.8 (0.55) | 0.16 (0.02) | **0.284** | **2.0 (0.1)** | **0.27 (0.04)** |
| 4 | 7.50 (0.32) | 0.61 (0.11) | 1 | 1 | 1 |

Table shows the mean (standard deviation across 20 segments) for the instantaneous frequency, amplitude, distance correlation, and frequency and amplitude ratios with base (IMF-4). In **bold** are shown values meeting harmonic conditions. Only IMF-3 passes all three tests and is a harmonic.

Hilbert transform into account, we added them to produce a single waveform, IMF-(3+4). This trace captured significantly more non-sinusoidality and showed the typical hippocampal theta waveform shape with a faster leading edge.

## V. DISCUSSION

Improvements in technology and analysis methods have shown the importance of non-sinusoidal waveform shape across species and modalities. Harmonics arising from non-sinusoidality have a significant impact on our analysis methods (e.g. on measures of phase-amplitude coupling) and hence impact our understanding of neural oscillations. Having precise vocabulary to describe these and test for them is thus only going to grow in importance. Our work provides much-needed clarity on what exactly a harmonic is. Our definition is based on rigorous, easy to understand conditions which match common intuitions about harmonics. We define two signals to be in a harmonic relationship if adding the higher frequency signal to the base does not change its period (equivalent to usual conditions of integer frequency ratios and a constant phase relationship) and if the joint oscillation has a well-defined instantaneous frequency (equivalent to saying there are no prominent secondary extrema.

Our work complements and contrasts with existing literature that deals with harmonics in neurophysiological data. As mentioned in the Introduction, most authors consider signals at an integer frequency ratio to the base (with a constant phase relationship) to automatically be a harmonic. The attempts to remove influence of harmonics from Fourier-based analyses have included identifying them using bicoherence [36] [8] and more recently by subtracting harmonic components from the signal [37]. Both of these are based around the Fourier spectrum. We propose that apart from this Fourier-transform understanding of harmonics, our understanding should also be informed by *the shape*. If a signal has large secondary extrema, it is no longer suitable to call it a single waveform.

Given our exploration of the above definition, we propose to understand harmonic structures as being of two types. The *weak* type has a well-defined instantaneous frequency (IF) for neighbouring harmonics, but harmonics introduce additional low-amplitude extrema and amplitude modulation leading to the IF being ill-defined in the limit of infinite harmonics. The *strong* type has a structure with a well-defined IF even for an infinite amount of harmonics and has no extrema beyond those of the base function. Some authors have previously suggested IF of the weak type is meaningless, and have proposed methods to restrict IMF bandwidth to disallow these [38]. Our distinction explains why this arises and distinguishes between the two harmonic structure types. Using an analytically tractable model and linking it to results about the Riemann Zeta function, we show why how the distinction arises. Weak harmonic structures are often the typical examples of harmonics outside Neuroscience (e.g. the saw-tooth function), so it makes sense to keep them included in the definition of harmonics whilst noting their difference to harmonics which do not introduce extrema. Interestingly, the harmonic identified in our LFP data was consistent with being of the strong type (*γ* exponent not significantly different from 2), as were harmonics in the FitzHugh-Nagumo model neuron (*γ* = 2.08). We postulate this is because real-world non-sinusoidal neural signals are derived from an underlying smooth variation in electrochemical properties. It is natural that resulting oscillatory waveforms are also smooth without new extrema, i.e. they are strongly harmonic. It qualitatively agrees with other known types of non-sinusoidal waveforms in the literature as well [1]. It would be interesting to explore the effect of simulated physiological parameters on the resulting harmonic structure and waveform shape.

Neural oscillations often come in bursts, i.e. include amplitude modulation (AM) [39]. Thanks to Bedrosian’s Theorem [17], all of the results in this paper apply to amplitude-modulated non-sinusoidal waveforms as long as the AM frequency is slower than that of the base function, making the spectra non-overlapping. We propose that talking about AM faster than the base function does not make sense anyway - the AM would be such that not even a single full cycle of the original waveform would be present. Thus, our results and conditions are fully applicable to sensible AM like that commonly present in neurophysiological data, as well as data that may change shape over time or across trials. In such non-stationary cases, our conditions should be applied dynamically to quasi-stationary or single trial epochs.

Empirical Mode Decomposition may complement and improve existing metrics based on the Fourier or Wavelet transform. This is because in theory, its IMFs can accommodate any non-sinusoidality. Instead of having to remove or worry about harmonic coupling, connectivity analyses can be performed directly on IMFs if all harmonics are present in one IMF. Our framework allows for a simple decision process for when to reconstruct highly non-sinusoidal waveform shapes in EMD analysis, where harmonics might be split across IMFs (Fig. 6). However, our approach relies on averaging across cycles to account for noise. As such, we sacrifice some of the single-cycle resolution due to noise. If single-cycle properties are being investigated, other forms of analysis might be more appropriate. These include a carefully designed manual mask, additional pre-processing, or using other EMD-based tools, e.g. iterated masking EMD [15]). Shape reconstruction (adding IMFs when appropriate) can also improve cases of poor sifts. Qualitatively, we have observed masked EMD to sometimes only partially extract the underlying base from the non-sinusoidal oscillation, leaving a waveform with negative instantaneous frequencies. Adding these IMFs reconstructs the underlying shape and makes EMD more robust to sifting details (e.g. mask amplitudes). Work on single-cycle waveform reconstruction in our group is ongoing.

The behaviour of summed tones and their instantaneous frequency can be explored by the readers with an interactive notebook accompanying this paper.

## VI CONCLUSION

Non-sinusoidal waveforms are ubiquitous in neurophysiology and our understanding of their clinical and functional importance is growing. The shape of neural oscillations has been previously shown to change with behaviour and disease (e.g. in Parkinson’s disease). Such waveforms are composed of sinusoidal harmonics present in the Fourier spectrum. However, a precise definition of when waveforms are to be considered harmonics is missing in the literature. In this work, we defined harmonic structures to be those that (i) have an integer frequency ratios and constant phases between constituent signals and (ii) have a well-defined (non-negative) instantaneous frequency. We showed this definition can be mathematically reformulated as specific conditions on the frequencies and amplitudes of the signals. We found two types of harmonic structures based on the presence of secondary extrema. We validated our framework on a simulated FitzHugh-Nagumo neuron and using EMD analysis of the hippocampal theta oscillation. In the latter, we showed why both conditions are important to unambiguously identify harmonics. Our work has important implications for metrics affected by non-sinusoidality, such as the phase locking value and other common measures of functional connectivity. Further work is needed to apply our framework in novel oscillation types and to explore the link between harmonic structure types and underlying generators of oscillations.

## APPENDIX

Interactive visualisations of the link between harmonics and instantaneous frequency are available *insert here*.

## A. Derivation of (5)

From basic notions of the Argand diagram, the phase angle *θ* is defined such that *θ* = arctan(*u/v*). The derivative of the inverse tangent is 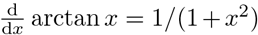. Hence, using the chain and quotient rules, we obtain the phase derivative as

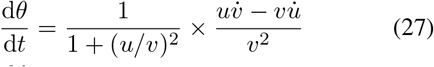

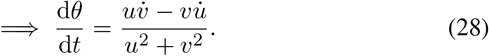

As the instantaneous frequency 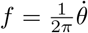, (5) follows.

## B. Derivation of (6)

We start with a general sum of sinusoids *u*, its Hilbert transform *v*, and their derivatives:

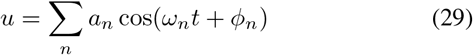

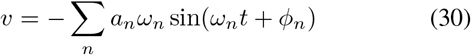

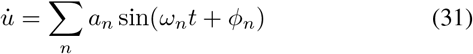

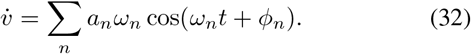

Putting these into the IF expression (5), combining the summations, and taking out a common factor of *a_n_a_m_ω_m_*, we find

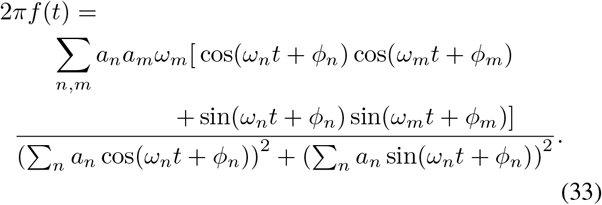

Applying the standard cosine difference formula in the numerator and omitting the denominator phase for legibility, we arrive at (6).

The terms in the numerator sum in (6) can be visualised as terms in an *N × N* matrix (again omitting the phase; ‘...’ indicates more elements along both axes):

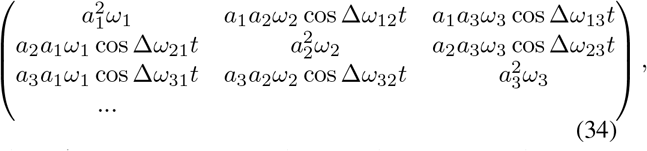

where Δ*ω_nm_* = *ω_n_ − ω_m_* and (33) is the sum of all the matrix elements. Demanding a well-defined joint IF *f _J_* ≥ 0 is thus equivalent to restricting the trace of (34) to be greater than the negative of the sum of all off-diagonal elements. It also makes it easier to see where (19) comes from, see below.

## C. Standard harmonic structures

In Fig. 8, we show the shape and instantaneous frequency of the triangular, saw-tooth, and square wave. Note how instantaneous frequency is well-defined except for delta function spikes at sharp edges. This means we are dealing with weak harmonic structures.

**Fig. 8.**
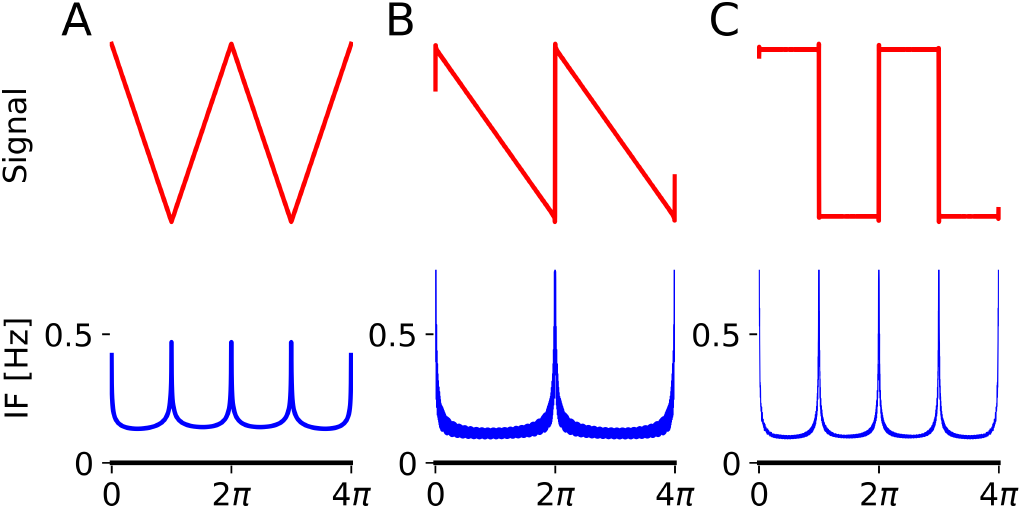
Standard harmonic structures. Top - signal and bottom - instantaneous frequency (IF) for (A) the triangular wave, (B) the saw-tooth wave, (C) the square wave (all formed from first 1000 harmonics). We see IF spikes near sharp edges but is otherwise well-defined.

## D. Derivation of (20)

Equation (19) can be trivially derived using the above matrix (34) by assuming *a*_1_ = 1, *a_n_* = 1/*n^γ^*, and *ω_n_* = *n*. Starting with (19) and applying the cosine double angle formula cos 2*x* = 2 cos^2^ *x* − 1, we get

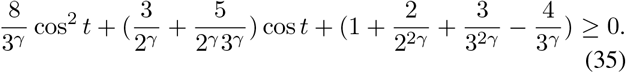

This is true so long the discriminant of the quadratic is negative, i.e. *b*^2^ − 4*ac* ≤ 0, where *a, b, c* are the coefficients in the quadratic. Thus, the critical exponent is when *b*^2^−4*ac* = 0:

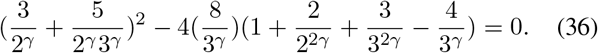

Expanding and simplifying this equation, we arrive at (20).

## E. Exponentially decaying harmonic structures

In the Main Text, we considered harmonic structures where the amplitude falls off as *a_n_* = 1/*n^γ^*. Here, we study another analytically tractable case of *a_n_* = exp(*λn*). As in the Main Text, consider the case with *ω_n_* = *n* and *ϕ_n_* = 0 and look at the maximum IF at *t* = 0. The numerator is

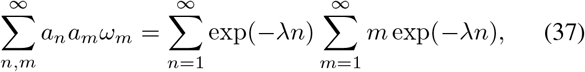

and the denominator simply 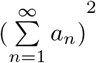. All these sums converge using standard geometric series methods provided *λ* > 0. Specifically,

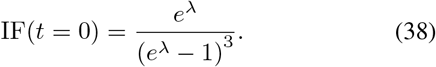

Harmonic structures with exponentially decaying amplitudes are thus always strongly harmonic. Interestingly, we found that in this case adding neighbouring harmonics can sometimes produce negative IF, which becomes non-negative again as the number of harmonics is increased. This shows how complex harmonics structures can be. However, the basic distinction based on presence of secondary extrema still holds.

